# Increased S6K1 phosphorylation protects against early steps of Tau aggregation under long-term mitochondrial stress

**DOI:** 10.1101/2022.06.21.496965

**Authors:** Lukasz Samluk, Piotr Ostapczuk, Dorota Malicka, Magdalena Dziembowska

**Affiliations:** Faculty of Biology, University of Warsaw, I. Miecznikowa 1, 02-096 Warsaw, Poland

**Keywords:** mitochondrial stress, mTOR pathway, S6K1 protein, protein aggregation, Tau protein, neurodegeneration

## Abstract

Many studies demonstrated the influence of mitochondrial stress on cytosolic signaling pathways. Here, we found that in cells upon long-term mitochondrial stress, phosphorylation of S6K1 protein, which is the mTOR pathway component, was increased, like in brains of Alzheimer’s disease (AD) patients. We checked if increased S6K1 phosphorylation was involved in Tau protein aggregation, which is one of AD hallmarks. HEK239T NDUFA11-deficient cells treatment with the mTOR inhibitor, INK128, or with S6K1 inhibitor, PF-4708671, caused the elevation of Tau aggregation. In contrast, stable overactivation of the mTOR pathway caused a further increase of S6K1 phosphorylation and reduced Tau oligomerization in HEK239T NDUFA11-deficient cells. Thus, we conclude that the increase in S6K1 phosphorylation is protective against Tau aggregation under mitochondrial stress.

## Introduction

Mitochondrial dysfunctions are considered as one of the causes of neurodegenerative diseases. Their potential involvement in the development of neurodegeneration is not only narrowed to the influence on energy metabolism. Accumulating data demonstrate that mitochondrial stress affects signaling pathways crucial for protein homeostasis and neuronal functions (Cabral-Costa and Kowaltowski, 2020). An example is the mTOR pathway which one of the numerous functions is the regulation of cellular protein synthesis via the mTORC1 complex (Fig. 1A). In our previous studies (Topf *et al*., 2018; Samluk *et al*., 2019) we have shown that dysfunctional mitochondria influence cellular protein homeostasis via the regulation of protein synthesis. Mitochondrial stress, accompanied by oxidative stress directly affected ribosomes by the oxidation of cysteine residues in ribosomal proteins (Topf *et al*., 2018) or by decreased phosphorylation of mTOR substrate, S6K1 protein, which resulted in dephosphorylation of S6 ribosomal protein (Samluk *et al*., 2019). It was demonstrated that the inhibition of cytosolic protein synthesis reduced mitochondrial degeneration (Wang *et al*., 2008) but prolonged induction of mechanisms that reduce translation under mitochondrial stress caused a loss of dendrites in neurons of *Drosophila melanogaster* (Tsuyama *et al*., 2017). Thus, to avoid cellular death, under long-term mitochondrial stress, protein synthesis needs to be restored. One of the mechanisms, which ensures a sufficient level of protein synthesis for cell survival under long-term mitochondrial stress is increased phosphorylation of S6K1 protein (Samluk *et al*., 2019). Interestingly, elevated S6K1 phosphorylation is also characteristic of the brains of Alzheimer’s disease patients, in which β-amyloid and Tau protein aggregates can be observed (An *et al*., 2003).

**Fig. 1.**
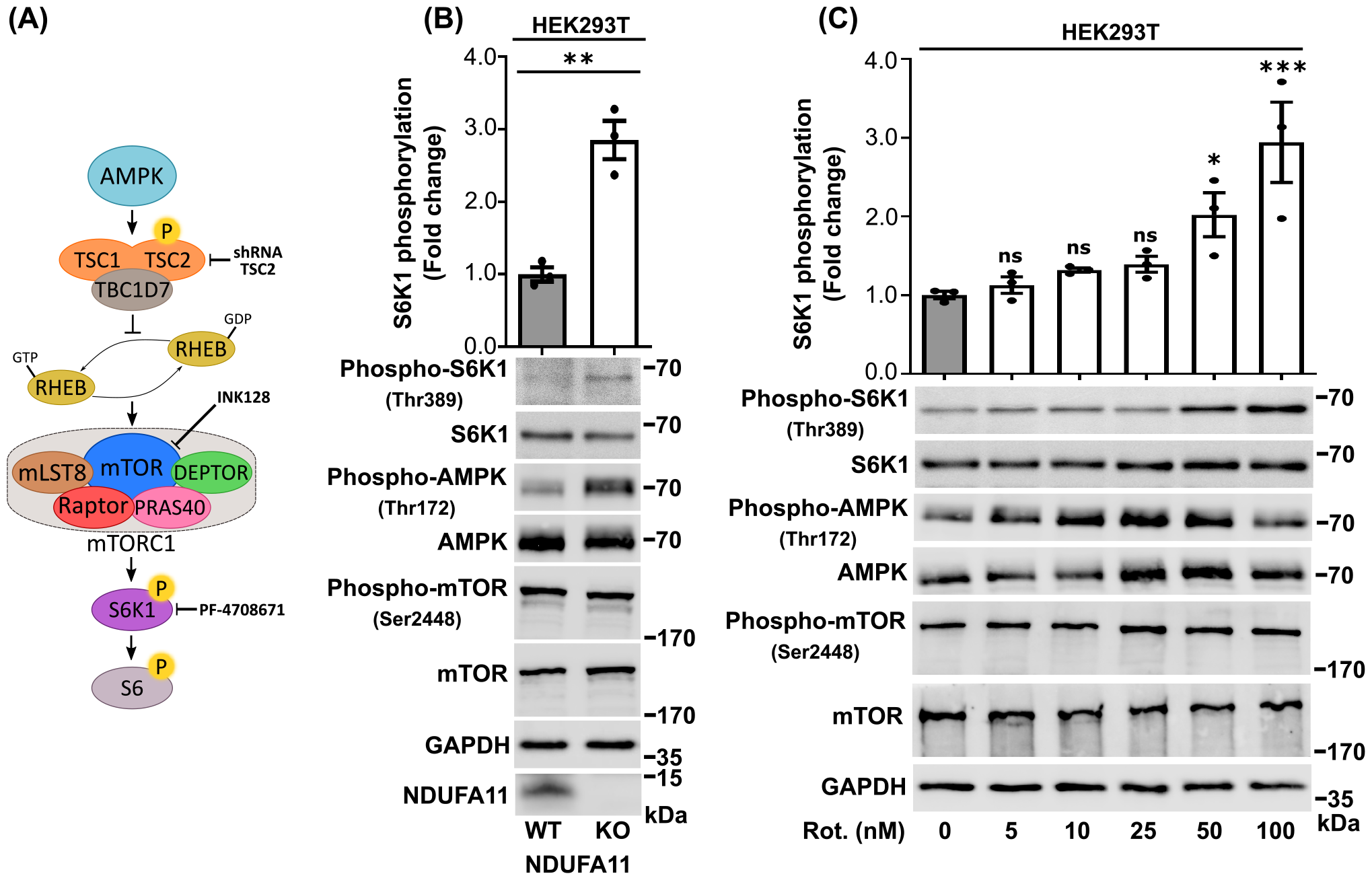
Long-term mitochondrial stress increases S6K1 (Thr389) phosphorylation. (A) Schematic diagram of mTORC1 signaling. Stress-activated AMPK phosphorylates TSC2 protein (in complex with TSC1 and TBC1D7), which inhibits RHEB (mTORC1 activator), resulting in a reduction of mTORC1 activity. The main components of mTORC1 are mTOR kinase, Raptor, DEPTOR, PRAS40, and mLST8 proteins. The dephosphorylation of S6K1 causes inhibition of its substrate (S6 ribosomal protein), resulting in a reduction of protein synthesis at the ribosome. INK128 is an mTOR kinase inhibitor, PF-4708671 is an S6K1 inhibitor, shRNA TSC2 was used for silencing of TSC2 resulting in TSC1-TSC2-TBC1D7 complex inhibition. (B) Phosphorylation of S6K1 (Thr389) in HEK293T wild type cells and in HEK293T NDUFA11 deficient cells. The data are expressed as mean ± SEM. n = 3. (C) Phosphorylation of S6K1 (Thr389) in HEK293T wild type cells that were treated for 48 h with rotenone. The data are expressed as mean ± SEM. n = 3. Rot, rotenone. ^*^p < 0.05; ^**^p < 0.01; ^***^p < 0.001; ns, not significant (p > 0.05) by Student’s t test (1B) or one-way ANOVA followed by Dunnett’s multiple comparisons test (1C).

A few recent studies demonstrated a direct impact of mitochondrial dysfunctions on the aggregation of proteins that are involved in the development of neurodegenerative diseases. The accumulation of mitochondrial precursor proteins in the cytosol as a result of mitochondrial protein import defects, induced the aggregation of α-synuclein and amyloid β (Nowicka *et al*., 2021a). Importantly, cytosolic protein aggregates were more efficiently cleared upon the stimulation of mitochondrial protein import (Nowicka *et al*., 2021b; Schlagowski *et al*., 2021). Moreover, it was shown that long-term mitochondrial stress induced early steps of Tau aggregation by increasing reactive oxygen species levels and affecting cellular proteostasis (Samluk *et al*., 2022). Interestingly, the increase in mitochondrial proteostasis by targeting mitochondrial translation and mitophagy reduced amyloid β aggregation in cells, worms and in transgenic mouse models of Alzheimer’s disease (Sorrentino *et al*., 2017). A previous study revealed that in mitochondrial prohibitin PHB2-deficient hippocampal neurons, Tau protein was hyperphosphorylated and aggregated but the mechanism of this phenomenon is still not known (Merkwirth *et al*., 2012).

In this study, we confirmed increased phosphorylation of S6K1 protein and enhanced Tau aggregation in mammalian cells under long-term mitochondrial stress. In order to monitor early steps of Tau aggregation, we performed the bimolecular fluorescence complementation (BiFC) assay using HEK293T cells that were treated with rotenone or HEK293T cells with TALEN-mediated knockout of gene encoding mitochondrial protein, NDUFA11 (Stroud *et al*., 2016). We demonstrated that mTOR and S6K1 inhibition with INK128 and PF-4708671, respectively, led to the escalation of Tau oligomerization in HEK293T NDUFA11-deficient cells. On the contrary, shRNA-mediated knockdown of TSC2 protein in HEK293T NDUFA11 knockout cells, which induced overactivation of the mTOR pathway and further increased S6K1 phosphorylation, caused the reduction of early steps of Tau aggregation. In the light of these findings, we consider increased phosphorylation of S6K1 as a beneficial adaptive response under long-term mitochondrial stress.

## Materials and methods

### Cell culture conditions

HEK293T cells were cultured in high-glucose (4.5 g/L) 90% Dulbecco’s Modified Eagle Medium (DMEM; Sigma, catalog no. D5671) supplemented with 10% FBS, 2 mM L-glutamine, 100 U/ml penicillin, 0.1 mg/ml streptomycin, and 50 μg/ml uridine at 37°C in a 5% CO_2_ humidified atmosphere. The cells were treated with rotenone (48 or 72 h) (Sigma, catalog no. R8875), mTOR kinase inhibitor (INK128, 3 h) (APExBIO, catalog no. MLN0128), and S6K1 inhibitor (PF-4708671, 24 h) (Sigma, catalog no. 559273) where indicated. HEK293T wild type and HEK293T NDUFA11 knockout cells were provided by David Stroud and Michael Ryan (Monash University, Melbourne, Australia) (Stroud *et al*., 2016). HEK293T NDUFA11 knockout cell line with shRNA-mediated knockdown of TSC2 was provided by Malgorzata Urbańska and Jacek Jaworski (International Institute of Molecular and Cell Biology, Poland) (Samluk *et al*., 2019).

### Cell culture transfection

HEK293T cells were seeded in 60 mm cell culture dishes and grown to reach 20-25% confluence on the day of transfection. For transfection of one plate, 6 µl of GeneJuice Transfection Reagent (Sigma, catalog no. 70967) was mixed thoroughly with 250 µl of Opti-MEM I Reduced Serum Medium (Gibco, catalog no. 31-985-070) and incubated for 5 min at room temperature. Next, purified plasmid DNA was added at a concentration of 0.5 or 1 µg/plate according to the specific experiment, gently mixed by pipetting and incubated for 15 min at room temperature. Then, GeneJuice/DNA mixture was added dropwise to each plate that contained 5 ml of complete high-glucose DMEM and gently rocked.

### Bimolecular fluorescence complementation

Bimolecular fluorescence complementation (BiFC) assay was performed to monitor the oligomerization of Tau protein (Fig. 2A) (Tak *et al*., 2013; Lim *et al*., 2014). HEK293T cells were transfected for 72 h with plasmids that encoded Tau protein that was fused to the N-terminal part of Venus protein, Tau-VN (VN-Tau (wt), Addgene, catalog no. 87368) and Tau protein that was fused to the C-terminal part of Venus protein, Tau-VC (Tau (wt) -VC, Addgene, catalog no. 873690) (Blum *et al*., 2015). The fluorescence of reconstituted Venus protein, which reflects Tau dimerization, was analyzed using a flow cytometer (BD LSRFortessa), a fluorescence microplate reader (Ex/Em 488/528) (Synergy H1 Hybrid Multi-Mode Microplate Reader, BioTek) and Zeiss LSM700 confocal microscope. Fluorescence was normalized to transfection efficiency that was verified by immunoblotting where indicated.

**Fig. 2.**
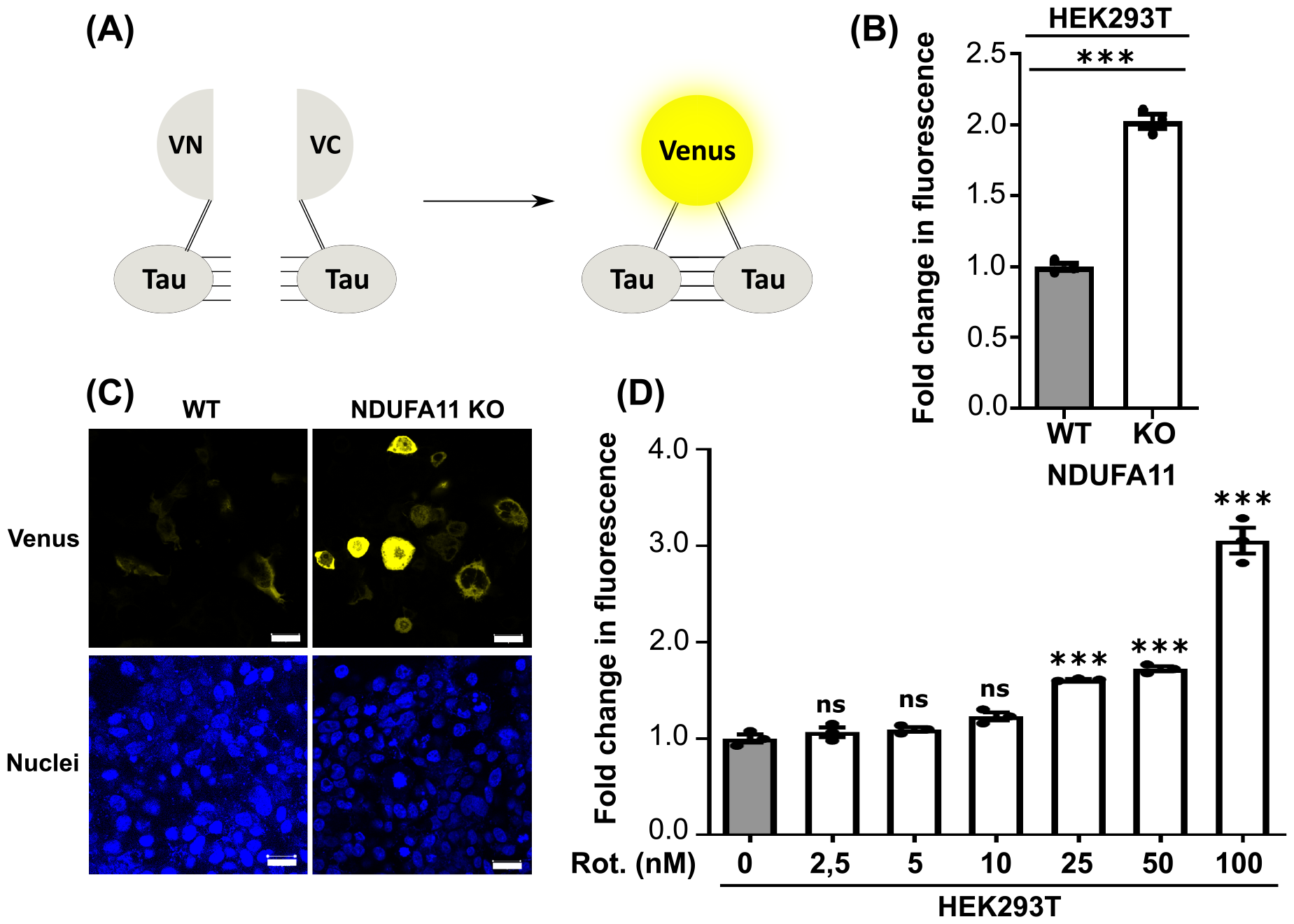
Tau protein aggregates in HEK293T cells with mitochondrial respiratory chain complex I dysfunction. (A) The principle of the bimolecular fluorescence complementation (BiFC) assay. Cells were transfected with plasmids that encoded Tau protein that was fused with the N-terminal part of Venus protein (Tau-VN) and Tau protein that was fused with the C-terminal part of Venus protein (Tau-VC). Tau protein aggregation resulted in the reconstitution of Venus protein and the increase in fluorescence. (B) Flow cytometry analysis of Venus protein fluorescence in HEK293T wild type cells (WT) and in HEK293T NDUFA11 knockout cell line. The data are expressed as mean ± SEM. n = 3. (C) Confocal images of Venus protein in HEK293T wild type (WT) cells and in HEK293T NDUFA11 deficient cells. Nuclei were stained with DAPI. Scale bar = 20 μM. (D) Flow cytometry analysis of Venus protein fluorescence in HEK293T cells that were treated for 72 h with rotenone. The data are expressed as mean ± SEM. n = 3. Rot, rotenone. ^***^p < 0.001; ns, not significant (p > 0.05) by Student’s t test (2B) or one-way ANOVA followed by Dunnett’s multiple comparisons test (2D).

### Immunofluorescence

HEK293T cells were transfected with BiFC plasmids for 72 h. The cells were then washed twice with PBS, fixed with 3.7% formaldehyde for 10 min at 4°C, washed again with PBS, and permeabilized for 5 min by treatment with 0.1% Triton X-100 in PBS. Then, cells were rinsed once with PBS and once with water. The samples were mounted in ProLong Diamond Antifade Mountant with DAPI (Thermo Fisher Scientific, catalog no. P36962) and analyzed with a confocal microscope (Zeiss LSM700).

### Miscellaneous

The RIPA buffer was used for the preparation of protein extracts. It contained 65 mM Tris-HCl (pH 7.4), 150 mM NaCl, 1% IGEPAL CA-630 (NP-40), 0.25% sodium deoxycholate, 1 mM EDTA, 2 mM PMSF (Sigma, catalog no. P7626), and phosphatase inhibitor cocktail (PhosSTOP, Roche, catalog no. 04 906 837 001). Proteins in Laemmli sample buffer that contained 50 mM dithiothreitol (DTT) were denatured at 65°C for 15 min. Total protein extracts were separated by SDS-PAGE on 12% gels. The following commercially available antibodies were used: S6K1 (Cell Signaling Technology, catalog no. 9202), Phopho-S6K1 (Thr389) (Cell Signaling Technology, catalog no. 9205), Phospho-S6 (Ser235/236) (Cell Signaling Technology, catalog no. 2211), S6 (Cell Signaling Technology, catalog no. 2217), Tau (TAU-5) (Merck, catalog no. 577801), GAPDH (Santa Cruz Biotechnology, catalog no. sc-47724), and TSC2 (Cell Signaling Technology, catalog no. 3612). Protein bands were visualized using secondary antibodies conjugated with horseradish peroxidase and chemiluminescence. Chemiluminescence signals were detected using Amersham Imager 600 RGB or x-ray films. Adobe Photoshop CS4 software was used for the digital processing of images and ImageJ software was used to quantify the immunoblots. The represented fold changes are means of fold changes that were obtained from independent biological replicates ± SEM. VN-Tau (wt) was a gift from Tiago Outeiro (Addgene plasmid no. 87368; http://n2t.net/addgene:87368; RRID:Addgene_87368; (Blum *et al*., 2015). Tau (wt)-VC was a gift from Tiago Outeiro (Addgene plasmid no. 87369; http://n2t.net/addgene:87369; RRID:Addgene_87369; (Blum *et al*., 2015). pRK5-EGFP-Tau AP was a gift from Karen Ashe (Addgene plasmid no. 46905; http://n2t.net/addgene:46905; RRID:Addgene_46905; (Hoover *et al*., 2010).

### Statistical analysis

Statistical Student’s t test or one-way ANOVA were performed using GraphPad Prism. Student’s t test or one-way ANOVA results are indicated consistently in all figures as ^*^ p < 0.05, ^**^p < 0.01, ^***^p < 0.001 and ns for not significant (p > 0.05).

## Results

### Long-term mitochondrial stress increase S6K1 phosphorylation

In our previous study we found that upon long-term mitochondrial dysfunctions phosphorylation of S6K1 was increased, contrary to short-term mitochondrial stress (Samluk *et al*., 2019). Interestingly, increased S6K1 protein phosphorylation was also observed in neurons treated for 6 h with mitochondrial stressors such as oligomycin and rotenone/antimycin-A, but this phenomenon was not explained (Zheng *et al*., 2016). Here, we confirm our previous observations that long-term mitochondrial stress is leading to the increase in the phosphorylation of S6K1 protein. HEK293T cells that were treated for 48 h with rotenone and HEK293T NDUFA11-deficient cells (impaired complex I-mediated respiration), exhibited enhanced phosphorylation of Thr389 residue of S6K1, as demonstrated by immunoblotting with the use of the antibody that recognizes this specific phospho-residue (Thr389) (Fig. 1B and C). In HEK293T NDUFA11 knockout cells the increase of S6K1 phosphorylation was observed despite the moderate reduction of the expression of this protein (Fig. 1B). In contrast, in HEK293T wild type cells expression of S6K1 protein was growing with the increase of applied rotenone concentration but anyway the increase of phosphorylation was much stronger than the increase of the S6K1 expression (Fig 1C). The highest S6K1 phosphorylation was observed in HEK293T wild type cells that received the highest dose of rotenone (100 nM) (Fig 1C), suggesting that the observed phenomenon is dependent on the intensity of mitochondrial stress. These results demonstrate that long-term mitochondrial stress lead to the increase of S6K1 phosphorylation.

### Reduction of the phosphorylation and activity of S6K1 under long-term mitochondrial stress leads to the increase of Tau aggregation

Previously, we have shown that enhanced phosphorylation of S6K1 caused increased phosphorylation of S6 ribosomal protein to stimulate ribosomes for protein synthesis under mitochondrial stress (Samluk *et al*., 2019). This adaptive response probably enabled sufficient protein synthesis and cell survival under these conditions but its effect on cellular proteostasis was unknown. Since the elevation of S6K1 activity was also observed in the brains of Alzheimer’s disease (AD) patients (An *et al*., 2003), we decided to check the influence of the increase of S6K1 phosphorylation on Tau protein aggregation. In order to monitor the early steps of Tau aggregation, we performed the bimolecular fluorescence complementation (BiFC) assay, which principle is shown in figure 2A. Tau protein dimerization and oligomerization were reflected by the increase in the fluorescence of Venus protein, which was detected using flow cytometry (Fig. 2B and D) and confocal microscopy (Fig. 2C). We confirmed our previous observations (Samluk *et al*., 2022) that Tau protein aggregation increased in HEK293T NDUFA11-deficient cells and HEK293T cells that were treated for 72 h with rotenone in a concentration-dependent manner (Fig. 2B, 2C and 2D). To determine the effect of the S6K1 phosphorylation on Tau aggregation we performed the BiFC assay in HEK293T NDUFA11 knockout cells that were treated for 3 h with INK128 (30 and 75 nM), which is an inhibitor of mTOR kinase (Fig. 3A). The phosphorylation of S6K1 was significantly reduced despite the increase in S6K1 expression upon cells treatment with INK128. Inhibition of mTOR activity under long-term mitochondrial stress resulted in the increase in the fluorescence that reflected Tau aggregation and it was dependent on increasing INK128 concentration (Fig. 3A). The results of fluorescence measurements were normalized to levels of Tau expression detected by immunoblotting to avoid the potential influence of unequal cell transfection. Next, in order to investigate the specific effect of S6K1 activity on Tau aggregation, BiFC assay was performed using HEK293T NDUFA11-deficient cells that were treated for 24 h with S6K1 inhibitor (20 and 50 µM), PF 4708671 (Fig. 3B). We observed dephosphorylation of S6 protein (Ser235/236), which is an S6K1 substrate, indicating that the activity of S6K1 was blocked. The normalized fluorescence was increased upon cells treatment with PF-4708671, showing that reduction of S6K1 activity led to the increase in Tau dimerization (Fig. 3B). These results demonstrated that inhibition of S6K1 phosphorylation and activity in NDUFA11 KO HEK293T cells triggered an increase in early steps of Tau aggregation.

**Fig. 3.**
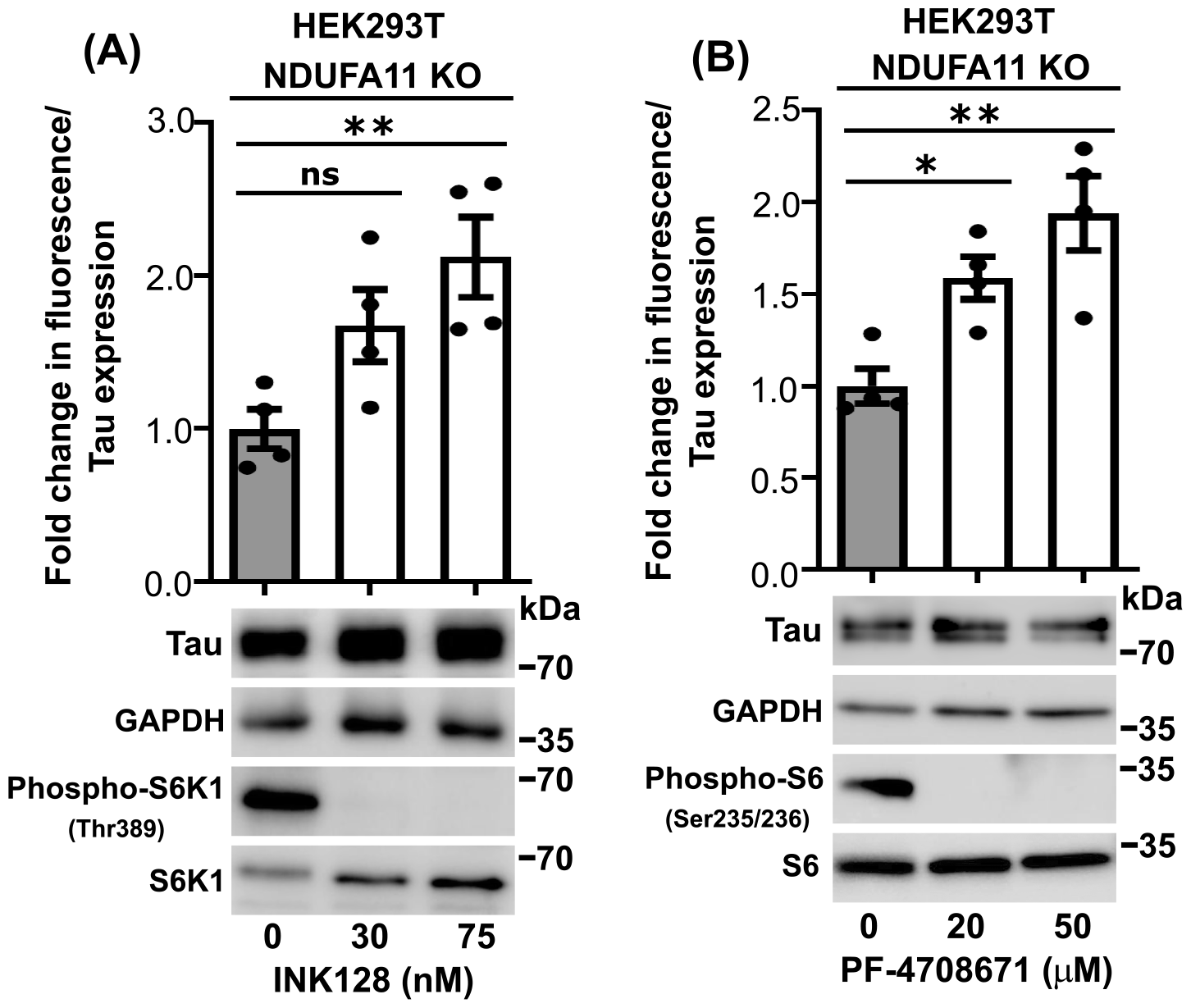
The inhibition of mTOR or S6K1 protein increased Tau aggregation in HEK239T NDUFA11-deficient cells. (A) Fold change in Venus fluorescence normalized to the level of Tau expression in HEK293T NDUFA11 knockout cells that were treated for 3 h with INK128 as indicated. The data are expressed as mean ± SEM. n = 4. (B) Fold change in Venus fluorescence normalized to the level of Tau expression in HEK293T NDUFA11 knockout cells that were treated for 24 h with PF-4708671 as indicated. The data are expressed as mean ± SEM. n = 4. ^*^p < 0.05; ^**^p < 0.01; ns, not significant (p > 0.05) (One-way ANOVA followed by Tukey’s multiple comparisons test).

### The activation of the mTOR pathway and the increase in the phosphorylation of S6K1 leads to the reduction of Tau aggregation under long-term mitochondrial stress

Next, we investigated whether activation of the mTOR pathway under conditions of long-term mitochondrial stress reduces Tau dimerization. Stable overactivation of the mTOR pathway was achieved by shRNA-mediated knockdown of TSC2 protein, which is a part of a protein complex that inhibits the mTOR pathway (Fig. 1A). As shown in figure 4A the level of TSC2 expression detected by immunoblotting was reduced approximately two times by specific shRNA compared to the control HEK293T NDUFA11-deficient cells. The knockdown of TSC2 led to the further increase of S6K1 phosphorylation at Thr389 and S6 protein at Ser235/236, suggesting overactivation of mTOR pathway (Fig. 4B). Next, we performed the BiFC assay. The fluorescence, normalized to Tau expression levels, was significantly decreased in HEK293T NDUFA11-deficient cells with a knockdown of TSC2 protein (Fig. 4B), indicating ower Tau dimerization under these conditions. Our results show that the activation mTOR pathway and increased S6K1 phosphorylation cause a significant reduction of early steps of Tau aggregation under long-term mitochondrial stress.

**Fig. 4.**
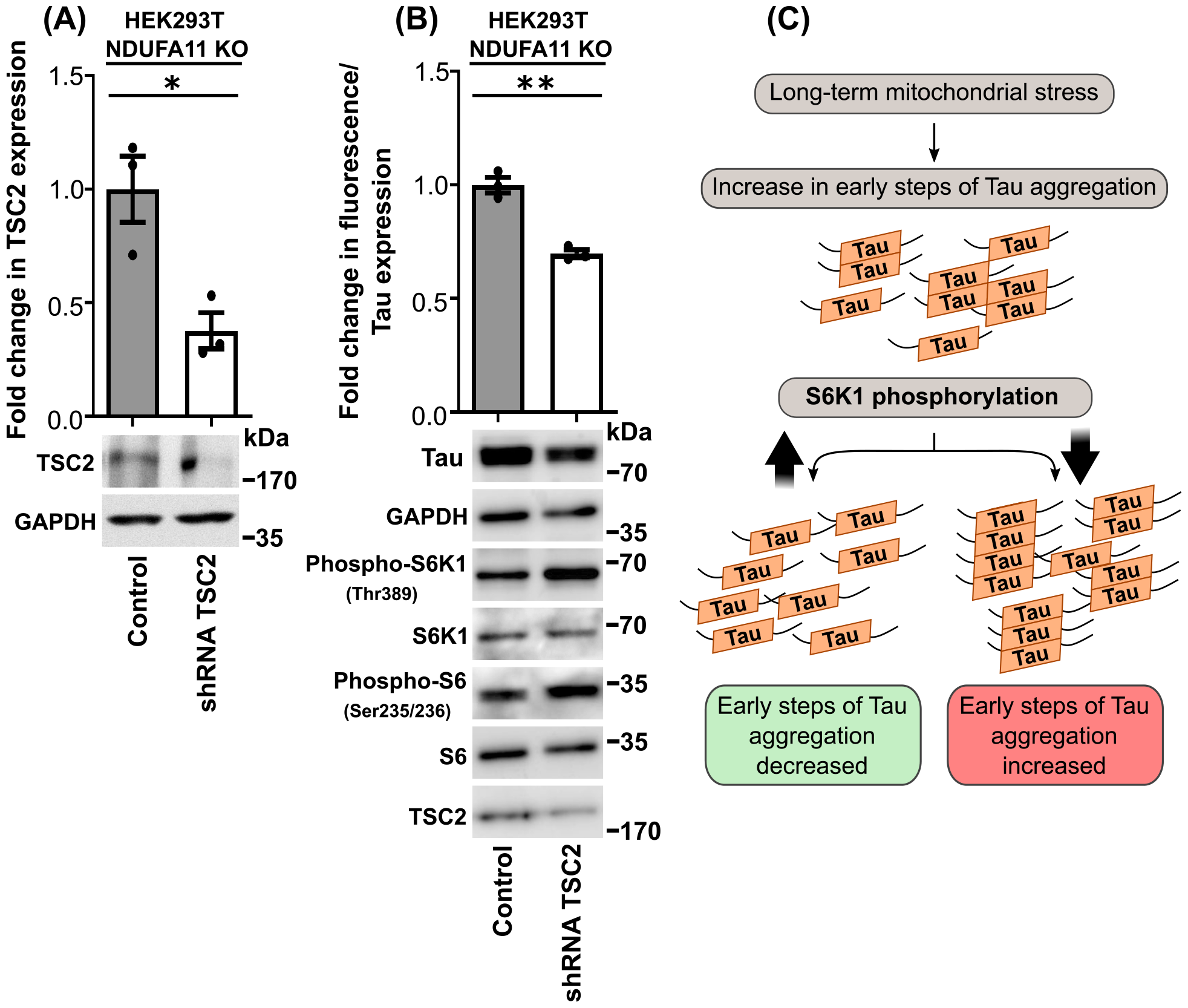
The knockdown of TSC2 protein reduced Tau aggregation in HEK239T NDUFA11-deficient cells. (A) Fold change in TSC2 expression in HEK293T NDUFA11 knockout cells with stable shRNA-mediated knockdown of TSC2 protein. The data are expressed as mean ± SEM. n = 3. (B) Fold change in Venus fluorescence normalized to the level of Tau expression in HEK293T NDUFA11 knockout cells with stable shRNA-mediated knockdown of TSC2 protein. The data are expressed as mean ± SEM. n = 3. (C) Schematic illustration of the influence of S6K1 phosphorylation on Tau protein aggregation under long-term mitochondrial stress. ^*^p < 0.05; ^**^p < 0.01 (Student’s t test).

## Discussion

Mitochondria, besides their primary function in energy production, are nowadays perceived as important signaling organelles. Under stress conditions, they affect signaling pathways that maintain protein homeostasis (proteostasis) in the cell. This fact puts mitochondria in a center of interest in studies investigating causes of neurodegenerative diseases, in which proteostasis defects and protein aggregation are observed. In the present and our previous study, we observed that long-term mitochondrial stress induce early steps of Tau aggregation (Samluk *et al*., 2022), characteristic of tauopathies, such as Alzheimer’s disease (AD). Previously we also demonstrated that mitochondrial dysfunctions affected the mTOR pathway (Samluk *et al*., 2019). This pathway among others regulates protein synthesis in the cell via phosphorylation of S6K1 protein that phosphorylates S6 ribosomal protein. Under the short-term mitochondrial stress phosphorylation of S6K1 was reduced, in contrast to long-term mitochondrial stress under which S6K1 phosphorylation was enhanced. Our interpretation of this phenomenon was that short-term inhibition of protein synthesis was beneficial for restoring protein homeostasis, while long-lasting protein synthesis inhibition may lead to the insufficient synthesis of crucial proteins and cellular death (Samluk *et al*., 2019). Thus, increased phosphorylation of S6K1 under long-term mitochondrial stress may be a pro-survival response but its effect on general protein homeostasis remains unknown. Two studies linked mitochondrial stress, increased S6K1 phosphorylation and neurodegeneration. The increase in S6K1 protein phosphorylation was observed in neurons that were treated for 6 h with compounds that affect mitochondrial function, oligomycin and rotenone/antimycin-A, (Zheng *et al*., 2016). Interestingly, the elevation of S6K1 activity was also observed in the brains of Alzheimer’s disease patients (An *et al*., 2003). In the present study, we studied the influence of the mTOR pathway activity and increased S6K1 phosphorylation on early steps of Tau aggregation. We demonstrated that general inhibition of the mTOR pathway and specific inhibition of S6K1 lead to the further increase in Tau aggregation under long-term mitochondrial stress. In contrast, overactivation of the mTOR pathway, which caused increased phosphorylation of S6K1 and its substrate S6 protein, significantly reduced Tau dimerization under long-lasting mitochondrial stress (Fig. 4C). These results suggested that the increase in S6K1 phosphorylation that was present in the brains of Alzheimer’s disease patients (An *et al*., 2003) could be a beneficial adaptive response. Moreover, our observations showed that further activation of S6K1 could be a method for the reduction of Tau aggregation. In contrast, the reduction of increased phosphorylation of S6K1 and S6 ribosomal protein, which was observed in mitochondrial myopathy, by mTOR inhibition caused the reversion of this mitochondrial disease (Khan *et al*., 2017). It seems that for the reduction of mitochondrial myopathy progression, inhibition of S6K1 and S6 phosphorylation which leads to the reduction of protein synthesis was more beneficial. In the case of tauopathies more beneficial was maintaining the synthesis of essential proteins for neuron functioning and the production of proteins that maintain proteostasis like molecular chaperons. All these results demonstrated that the mTOR pathway is an important player in mitochondrial and neurodegenerative diseases but in order to obtain a therapeutic effect of pharmacological interventions, i.e. activation versus inhibition of the mTOR pathway, the treatment need to be tailored to the particular pathology.

## Abbreviations

AD: Alzheimer’s disease
AMPK: adenosine monophosphate-activated protein kinase
BiFC: bimolecular fluorescence complementation
DEPTOR: DEP domain-containing mTOR-interacting protein
DMEM: Dulbecco’s Modified Eagle Medium
DTT: dithiothreitol
EDTA: ethylenediaminetetraacetic acid
GAPDH: glyceraldehyde-3-phosphate dehydrogenase
mLST8: mammalian lethal with SEC thirteen 8
mTOR: mechanistic target of rapamycin
mTORC1: mTOR complex 1
NDUFA11: NADH:ubiquinone oxidoreductase subunit A11
PBS: phosphate-buffered saline
PMSF: phenylmethylsulfonyl fluoride
PRAS40: proline-rich AKT substrate of 40 kDa
Raptor: regulatory-associated protein of mTOR
S6K1: ribosomal protein S6 kinase 1
SDS-PAGE: sodium dodecyl sulfate-polyacrylamide gel electrophoresis
SEM: standard error of the mean
TSC2: tuberous sclerosis complex 2

## Acknowledgments

We thank David Stroud and Michael Ryan for providing the HEK293T NDUFA11 knockout cell line, Malgorzata Urbańska and Jacek Jaworski for providing the HEK293T NDUFA11 knockout cell line with shRNA-mediated knockdown of TSC2 and Minji Kim and Agnieszka Chacińska for valuable comments and discussions. This work was supported by National Science Centre of Poland (NCN), Grant Number 2019/33/B/NZ3/00533 (for LS).

## Author contributions

LS conceived, designed, and supervised the study. LS, PO and DM performed the experiments and analyzed the data. LS wrote the manuscript with the input of PO and MD.

